# Addiction-Related Memory Transfer and Retention in Planaria

**DOI:** 10.1101/2021.09.12.459965

**Authors:** Kenneth Samuel, Easter S. Suviseshamuthu, Maria E. Fichera

## Abstract

Memory retention and transfer in organisms happen at either the neural or genetic level. In humans, addictive behavior is known to pass from parents to offspring. In flatworm planaria (*Dugesia tigrina*), memory transfer has been claimed to be horizontal, i.e., through cannibalism. Our study is a preliminary step to understand the mechanisms underlying the transfer of addictive behavior to offspring. Since the neural and neurochemical responses of planaria share similarities with humans, it is possible to induce addictions and get predictable behavioral responses. Addiction can be induced in planaria, and decapitation will reveal if the addictive memories are solely stored in the brain. The primary objective was to test the hypothesis that addictive memory is also retained in the brainless posterior region of planaria. The surface preference of the planaria was first determined between smooth and rough surfaces. Through Pavlovian conditioning, the preferred surface was paired with water and the unpreferred surface with sucrose. After the planaria were trained and addicted, their surface preference shifted as a conditioned place preference (CPP) was established. When decapitated, the regenerated segment from the anterior part containing the brain retained the addiction, thus maintaining a shift in the surface preference. Importantly, we observed that the posterior part preserved this CPP as well, suggesting that memory retention is not attributed exclusively to the brain but might also occur at the genetic level. As a secondary objective, the effects of neurotransmitter blocking agents in preventing addiction were studied by administering a D1 dopamine antagonist to planaria, which could provide pointers to treat addictions in humans.

## 1 Introduction

Planaria are flatworms from the kingdom Animalia, of the phylum Platyhelminthes, class Rhabditophora, and in the order Tricladida (Buttarelli et al., 2008). They include several species of flatworms that have bilateral symmetry, are triploblastic, and acoelomate organisms (Cebrià, 2016). These invertebrates are commonly present in aquatic habitats, the majority of them in freshwater bodies like ponds, lakes, and rivers with some exception in saltwater and terrestrial environments (Justine et al., 2018). Planaria show a preference toward electric fields, magnetic fields, and chemical gradients (Shomrat & Levin, 2013). They are sensitive to light and prefer darker environments (Inoue et al., 2004). Similarly, they have been shown to prefer different surface textures, rough over smooth (Jawad et al., 2017). Planaria have the ability to reproduce asexually via binary fission (Hagstrom et al., 2016). A fragment of planaria can regrow into a completely new organism (Cebrià, 2016). This is due to the high concentration of stem cells, around 25%, which are pluripotent neoblasts (Hammoudi et al., 2018). Once a worm is divided into two or more sections, each section develops into an independent organism but remains as a genetic clone (Deochand et al., 2018).

Planaria have a very high regenerative capacity including their nervous system development. They are easily accessible, cost-effective, easy to maintain, and do not require any special equipment, making them a good choice for lab experiments. They exhibit specific behavioral responses to psychoactive substances. Therefore, they are used as a model organism in various studies that investigate chemical responses, behavioral pharmacology, and toxicology (Pagán et al., 2009). On the other hand, studies on neurotoxicology rely heavily on *in vivo* animals such as rodents since they have evolutionary proximity to humans. But these mammalian model systems are often expensive and time-consuming to support and may present ethical issues when using environmental toxicants.

In planaria, the head located in the anterior part houses a primitive “brain,” which contains specialized sensory organs and nerve cells (Pagan, 2019); Buttarelli et al., 2008). A closer observation at the neurological and neurochemical levels reveals many similarities between planaria and vertebrates. The nervous system of planaria has several resemblances to vertebrates, including cell morphology and physiology (Pagán et al., 2009). Planaria possess almost every neurotransmitter found in mammals, including dopamine (Buttarelli et al., 2008). Moreover, the neurotransmitters that are used to communicate information between different nerve cells and their receptors in planaria are not too different from those of humans. The planarian brain can be described as bilobed and has various neural characteristics analogous to the human brain (Sarnat & Netsky, 1985). Therefore, many studies have been conducted on planaria’ s physiological and regenerative properties by manipulating their physical and chemical environments (Hagstrom et al., 2016). Due to similar neuromuscular communication methods, conclusions on relevant mechanisms in humans can be determined from mechanistic studies in planaria. Furthermore, the amputated planaria quickly grow back their missing parts (Cebrià, 2016) enabling behavioral assays to be carried out on growing and developed adults in parallel, and observations to be made regarding toxicity to the developing brain (Hagstrom et al., 2016). With these unique features including neuronal complexity and neurotransmitters similar to humans, studies on the behavioral response following exposure to different drugs are possible with planaria instead of mammals or humans (Buttarelli et al., 2008).

Research has shown that there are some genetic factors behind addictions. Some addiction traits might be passed down from parents to offspring, but this remains a grey area. Irrespective of the abusive substance, about 40-60% of the addictive risk in humans is genetic (Szalavitz, 2015). For example, children of alcohol abusers tend to get addicted 3-5 times more likely than a child with non-alcoholic parents (Szalavitz, 2015). This might be caused by epigenetic mechanisms where the activity of genes can be switched on or off. For instance, cocaine binging can cause the expression of certain genes that are normally in a state of dormancy in adults, and hence rewires the brain (Szalavitz, 2015). A suitable organism to test if addictions are genetic would be the planaria since they have a complex neural network as well as cephalic structures and neurotransmitters similar to vertebrates. Planaria are used to study drug abuse and behavioral responses (Pagán, 2017), as they are capable of acquiring drug-induced addiction behaviors similar to vertebrates. Sufficient drug exposure can develop physical dependence and induce withdrawal responses in planaria—drugs like cocaine, nicotine, opioids, amphetamines, and cannabinoids produce abstinence-induced withdrawal symptoms (Pagán, 2017).

Sucrose has also been shown to elicit addictive behaviors, and the neurochemical response to binging on sucrose is similar to that of addictive drugs (Zhang et al., 2013). Sucrose has been characterized as a drug of abuse in animals and its regular exposure provokes behavioral and physiological responses in animals. Planaria exposed to a solution of 1-10% sucrose develops a conditioned place preference (CPP). This can be observed by using the Pavlovian response where contextual cues are paired up with rewarding agents e.g., sugar (Jawad et al., 2017).

Another related research is understanding learning abilities and memory retention in planaria due to the organization of cranial ganglia. Conditioning has been achieved in these flatworms and they are known to retain behaviors that were learned (Abbott & Wong, 2008). Planaria were shown to retain long-term memory, enough to persist until the regeneration of a new brain after decapitation (Shomrat & Levin, 2013). If a trained planaria is consumed by an untrained planaria, learning could be transferred from the trained to the untrained one. However, controversies surround the memory transfer in planaria through cannibalism. These experiments were predominantly conducted by McConnell (1962) but were written off as myths and forgotten (Deochand et al., 2018). A few years later, another study suggested that memory could be transferred interspecifically when a wild planaria consumed a conditioned one (Ragland & Ragland, 1965). Recent molecular experiments examining RNA interference support the idea of memory being transferred through cannibalism (Deochand et al., 2018). A research question yet to be validated is whether the memory stored in each genetic clone of planaria is identical after decapitation since one segment will contain the brain and the other will not.

The primary aim of our research was to observe if the addictive memory is retained in the anterior and, more specifically, the posterior segment of planaria after they undergo a dissection. If an adult planaria is induced into an addiction to a rewarding agent, e.g., sucrose, certain genes will be expressed. The addiction can be observed by their behavior through learned Pavlovian conditioned responses. We can verify if the addiction has been passed on to its offspring by dissecting them since they are capable of reproducing by binary fission. The brain will be absent in the posterior part, which will develop into a new organism. If addiction is caused by the expression of certain genes, the brainless segment would still retain the addictive memory. We hypothesized that the brainless posterior segment will retain the addictive memory even after the head containing the brain has been removed.

Drug abuse destabilizes the amount of some neurotransmitters including a rise in the extracellular concentration of dopamine (Volkow et al., 2009). The dopamine level in planaria is directly related to its motility and movement control (Buttarelli et al., 2000), thereby resulting in higher activity levels (Pagán et al., 2009). D1 and D2 dopamine antagonists have been shown to obstruct sucrose addictions (Zhang et al., 2013), as they fix to different dopamine receptors and disrupt the fixation of dopamine with those receptors. As a secondary aim, we tested if adding a D1 dopamine antagonist to the sugar solution helps inhibit the CPP for the rewarding agent. The D1 antagonist was chosen since D1 receptors are the most abundant dopamine receptors found in the central nervous system (Bhatia et al., 2020). We hypothesized that the presence of a D1 dopamine antagonist will prevent the planaria from getting addicted to the sugar solution. Experiments measuring the distance covered by planaria exposed to the D1 dopamine antagonist have proven to inhibit a CPP formation (Jawad et al., 2017). However, our experiments conducted with the antagonist focused on the surface preference of planaria by observing the time spent on different surfaces.

First, we verified whether both the anterior and posterior fragments resulting from decapitation prefer rough surfaces since planaria inherently maintain this surface preference as reported in (Jawad et al., 2017). Second, we investigated if the memory was stored in the posterior segment by decapitating the planaria after a CPP was established. Third, the planaria were treated with a D1 dopamine antagonist to find if the addiction could be prevented by the antagonist.

## 2 Materials and Methods

### 2.1 Subjects

We performed the experiments with 52 planaria specimens that belong to the following two species: brown planaria (*Dugesia tigrina*) and black planaria (*Dugesia dorotocephala*). The planaria were kept in a bowl containing Evian natural spring water filled up to 1 cm, at room temperature. They were fed egg yolk before the commencement of the experiment. Feeding was stopped a day before starting the training period.

### 2.2 Materials

Three Petri dishes of 9 cm diameter were modified as described in the next section. In order to create a rough surface, we used white sand and transparent silicone glue (DAP 00688 all-purpose adhesive sealant, 100% silicone, 2.8-ounce tube). Sucrose was needed to prepare a sugar solution, and a D1 dopamine antagonist (SCH-23390 hydrochloride, Sigma Aldrich) was obtained to test the secondary hypothesis. The motility of the planaria was measured using 0.5 cm^2^ graph paper.

### 2.3 Methods

#### 2.3.1 Preparation of the Petri Dishes and Surfaces

One Petri dish was kept unaltered to serve as a smooth surface for the planaria to glide on (left panel of Fig. 2). White sand was stuck to the inner and bottom surface of the second dish to make it rough (right panel of Fig. 2). The silicone adhesive used for this purpose took about 72 hours to dry completely and cure properly. The third Petri dish was divided into two sections as follows: one half was left smooth while the other half was glued with sand (hereafter referred to as half-rough dish, middle dish shown in Fig. 2). We prepared a one-liter solution containing 10% sucrose for training the planaria.

**Figure 1:**
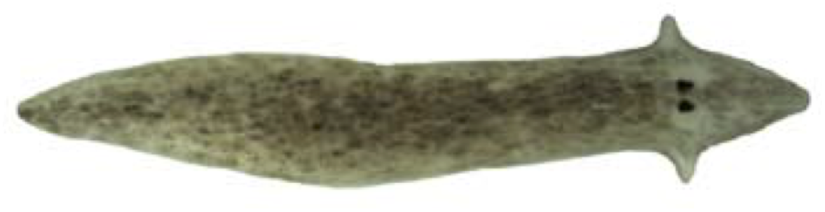
A specimen of *Dugesia dorotocephala* investigated in this study. Figure courtesy of Accorsi et al., 2017.

**Figure 2:**
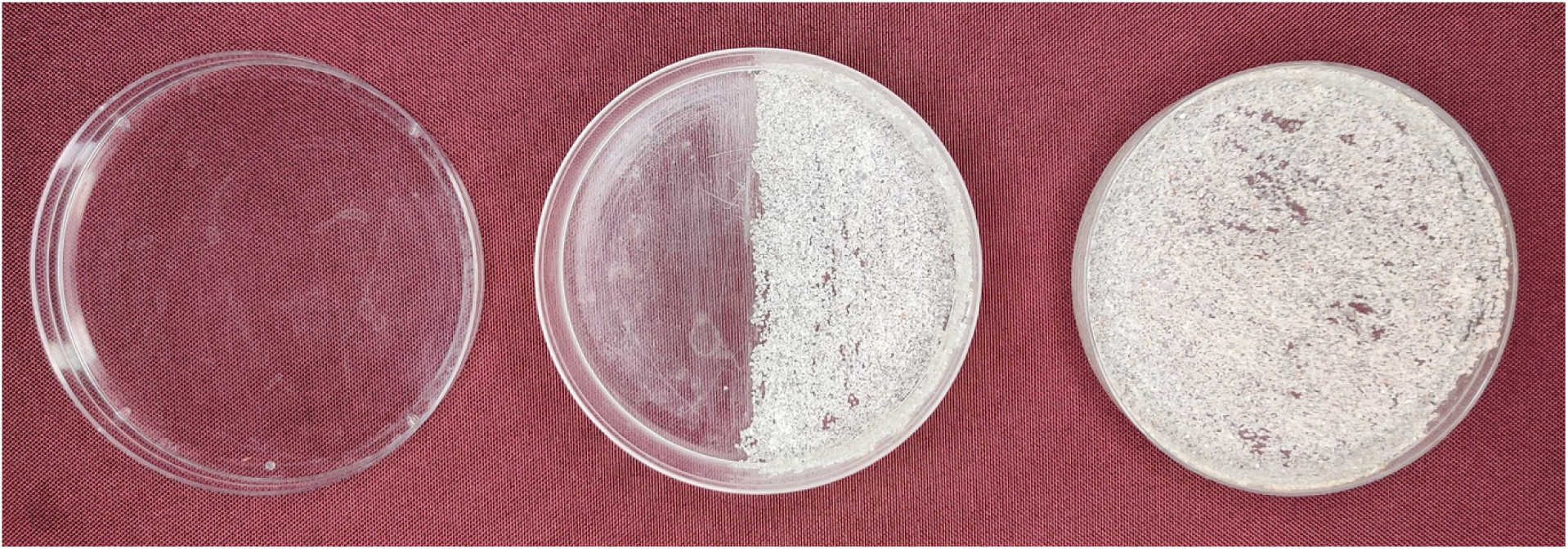
Training phase Petri dishes. Left: smooth surface, paired with sugar solution. Middle: half-rough dish, divided into rough and smooth halves to be used for the pre-and post-training tests. Right: rough surface paired with water.

#### 2.3.2 Motility in sugar or water solution

The behavior of planaria in sugar solution and water was assessed by observing its motility by counting the number of gridline crosses. A planaria was observed by placing it in water followed by the sugar solution for eight minutes. The planaria was kept in a 6 cm diameter Petri dish which was placed over a 5 mm^2^ graph sheet. Its movement was determined by counting the number of times the head crossed a gridline of the graph sheet.

#### 2.3.3 Establishing Conditioned Place Preference

The experiment was divided into the following three phases.

Phase 1 – Pre-training Phase: On the first day, 15 planaria were transferred to the half-rough dish for 30 minutes. The time spent by each planaria on either surface (smooth and rough) was recorded. The surface preferred by a planaria was determined to be the one where it spent a longer time. Thus, the duration of stay on a surface was considered to be the indicator for the level of preference.

Phase 2 – Training Phase: The preferred surface was then paired with water (preferred+water) and the unpreferred surface with 10% sugar solution (unpreferred+sugar). The training phase was designed to last for 12 days (30 minutes each day). During this period, the planaria were alternated between the preferred+water and unpreferred+sugar surfaces.

Phase 3 – Post-training Phase: After 12 days, the planaria were tested to see if their surface preference had shifted. Each individual was placed on the half-rough Petri dish for 30 minutes and the respective time spent on the rough and smooth surface was noted.

#### 2.3.4 Surface Preference of Untrained Decapitated Planaria

Ten planaria were amputated into head and tail segments, which were then placed on the half-rough dish for 30 minutes. The number of segments on each surface was counted every minute and recorded.

#### 2.3.5 Verifying Memory Retention

The establishment of a CPP would lead the planaria to opt for the unpreferred surface, thus associating the preference to the presence of sugar. To verify whether the memory is retained at a genetic level, the planaria were amputated into two fragments, and the post-training phase was repeated for the individuals regenerated from the anterior and posterior regions. The time spent by the regenerated planaria on the preferred surface was then recorded.

#### 2.3.6 Investigating Effects of a Dopamine Antagonist

This investigation was carried out by adding 1-μM of dopamine antagonist to the 10% sugar solution (unpreferred+sugar+antagonist). The training and the post-training phases were repeated. The training phase consisted of alternating between the preferred+water and unpreferred+sugar+antagonist solution for 30 minutes each day, for 12 days. The post-training phase was also repeated for 30 minutes on the half-rough dish in order to verify the retention of the planarian sucrose addiction in the presence of dopamine antagonist by observing the surface preference of the planaria.

## 3 Results

### 3.1 Behavior in Water and Sugar Solution

The gridline crossings demonstrated that the planaria displayed a higher degree of motility in the sugar solution than water. The cumulative gridline crossings by the planaria for eight minutes in water and the sugar solution are shown in Fig. 3.

**Figure 3:**
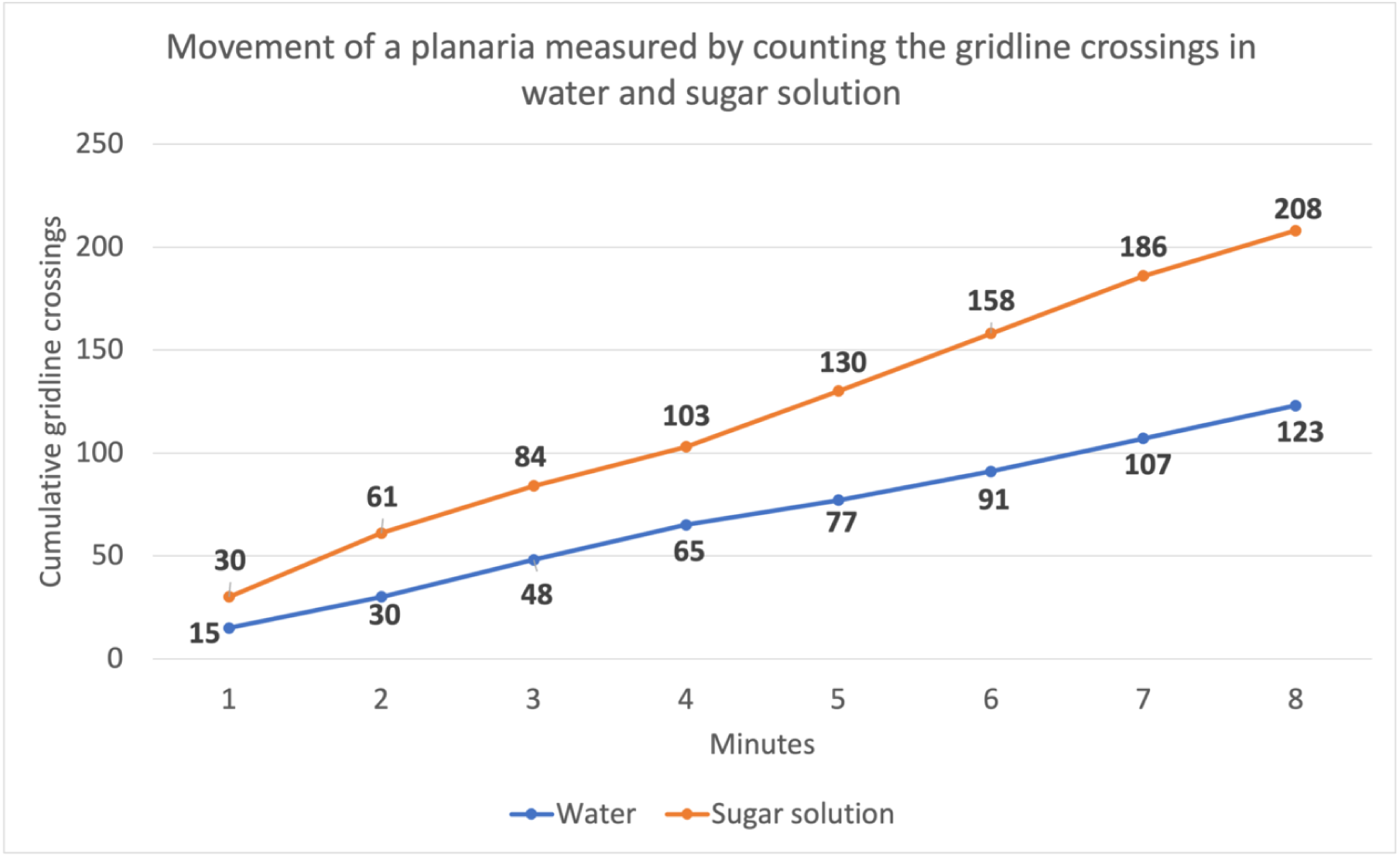
The cumulative number of times the head of a planaria crossed the gridlines in a graph sheet, when the planaria was placed in water (blue) and a sugar solution (orange).

### 3.2 Surface Preference of Untrained Planaria

This experiment was intended to identify the preferred surface of each of the 15 untrained planaria. For this purpose, the planaria were individually placed on the half-rough dish for 30 minutes. Then we recorded separately the time (in seconds) spent by each planaria on the rough surface and the smooth surface in this duration. As depicted in Fig. 4, the planaria spent an average time of 1025.53 seconds on the rough surface and 774.47 seconds on the smooth surface, thus showing a preference for the rough over the smooth surface.

**Figure 4:**
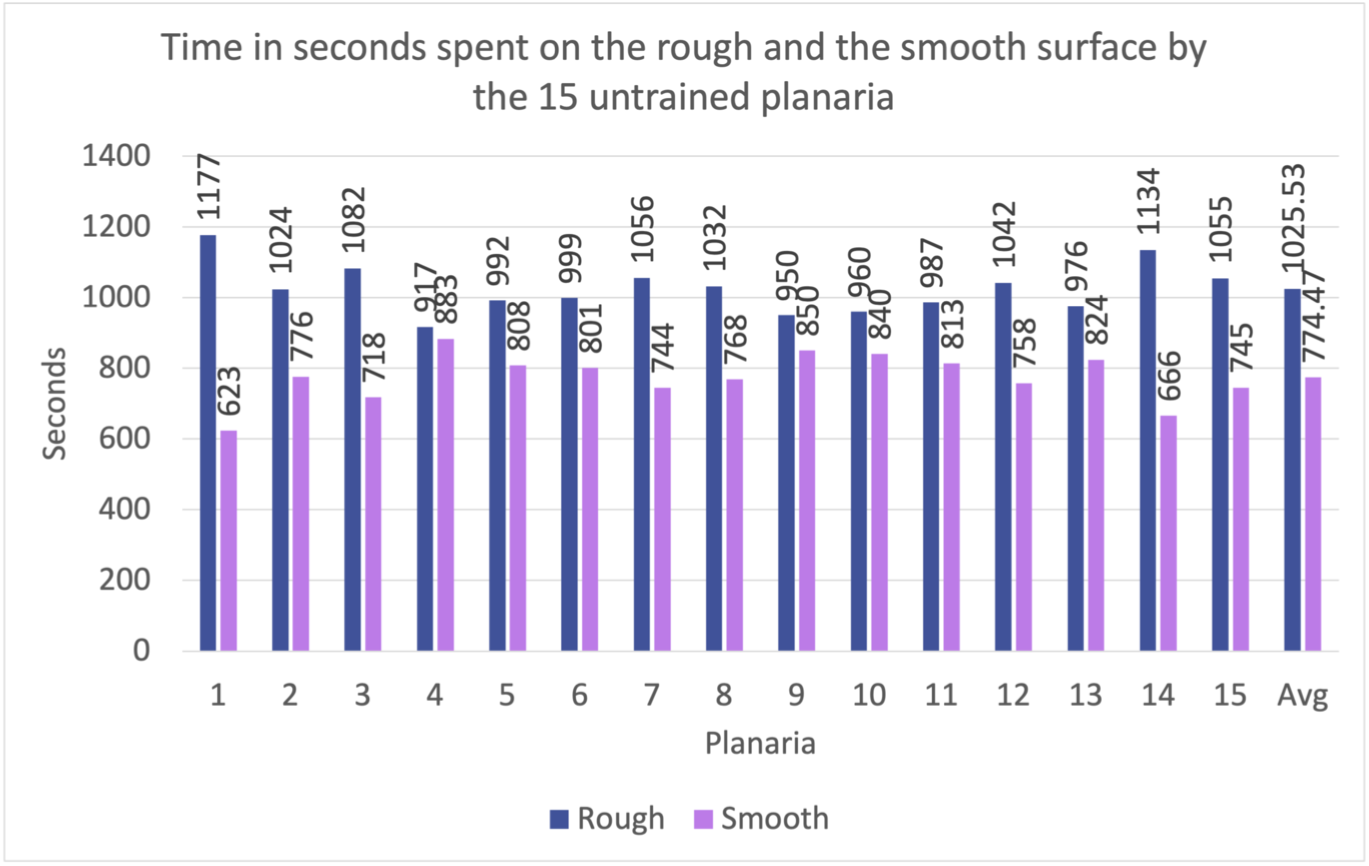
Surface preference of the 15 untrained planaria. The rough surface is preferred by all the untrained planaria as evidenced by the longer time spent by each planaria on the rough surface (blue) than on the smooth surface (purple).

### 3.3 Surface Preference of Trained Planaria

Once the surface preference of the planaria was established, the training phase commenced by pairing the rough (preferred) surface (R) with water (W), referred to as R+W, and pairing the smooth surface (S) with 10% sugar solution (S), denoted as S+S. For the next 12 days, the planaria were exposed to the R+W and S+S alternatively every day for 30 minutes. On the 13^th^ day, the post-training phase began. Each planaria was individually transferred back to the half-rough dish for 30 minutes and the time spent on the rough and smooth surface was recorded as in the case of untrained planaria. We found that the planaria spent an average of 607 seconds on the rough surface and 1193 seconds on the smooth surface. Fig. 5 shows that the preference of all the experimented planaria had shifted to the smooth surface after the training.

**Figure 5:**
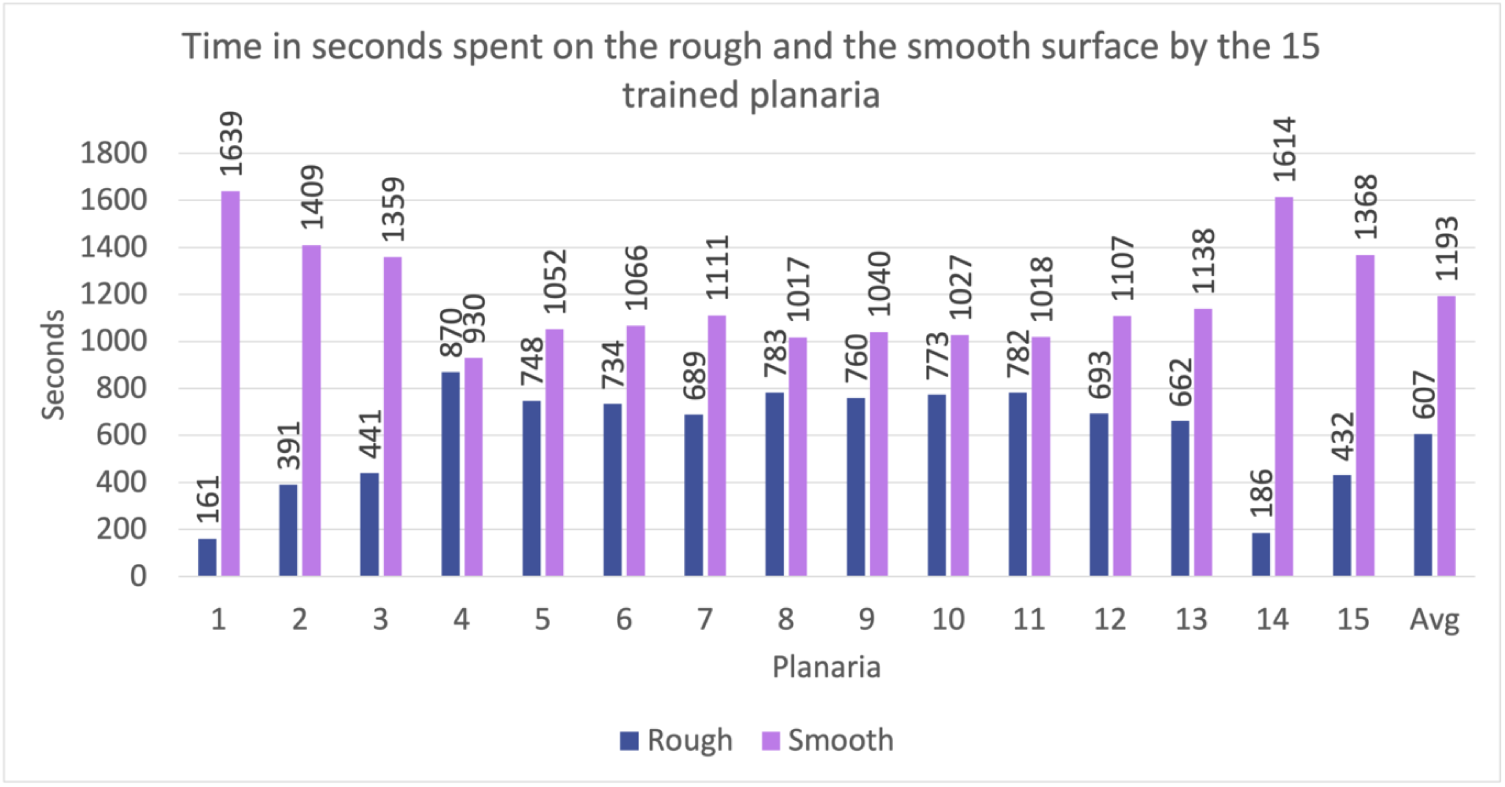
Surface preference of the trained planaria. The smooth surface is preferred by the trained planaria as the time spent on this surface (purple) is longer compared to that on the rough surface (blue).

### 3.4 Surface Preference of Decapitated Planaria

Fifteen trained planaria were further trained for another four days. After this mini-training phase, they were decapitated. All the 30 segments, i.e.,15 anterior segments containing the head and the remaining posterior ones without the head, were transferred to the rough side (unpreferred surface of the trained planaria) of the half-rough dish. The surface preference was determined by counting the number of segments on each surface every minute for 30 minutes.

To compare the preference of the decapitated trained planaria segments with untrained segments, 10 untrained organisms were also amputated into head and tail fragments. The resulting 20 head and tail segments were placed on the smooth side (unpreferred surface of the untrained planaria) of the half-rough dish, and the number of segments on each surface was noted every minute for 30 minutes.

Of the trained decapitated planaria, both the anterior and posterior segments showed a preference for the smooth surface and began to move toward the smooth surface progressively. By the end of 30 minutes, 27 segments had transitioned to the smooth surface while only three segments remained on the rough surface as depicted in Fig. 6(left). As for the untrained segments, there was a gradual movement toward the preferred rough surface. After 30 minutes, 16 segments were present on the rough surface and the remaining four on the smooth surface as shown in Fig. 6(right). Interestingly, the rate at which the transition took place toward the preferred surface is higher in the case of trained planaria compared to the untrained planaria. The transitions were mathematically modeled by the exponential decay curves in Fig. 7 for the trained planaria in blue and the untrained planaria in purple. The slope of the blue curve representing the transition rate of trained planaria is 0.20, whereas the slope of the purple curve denoting the transition rate of the untrained planaria is 0.07.

**Figure 6:**
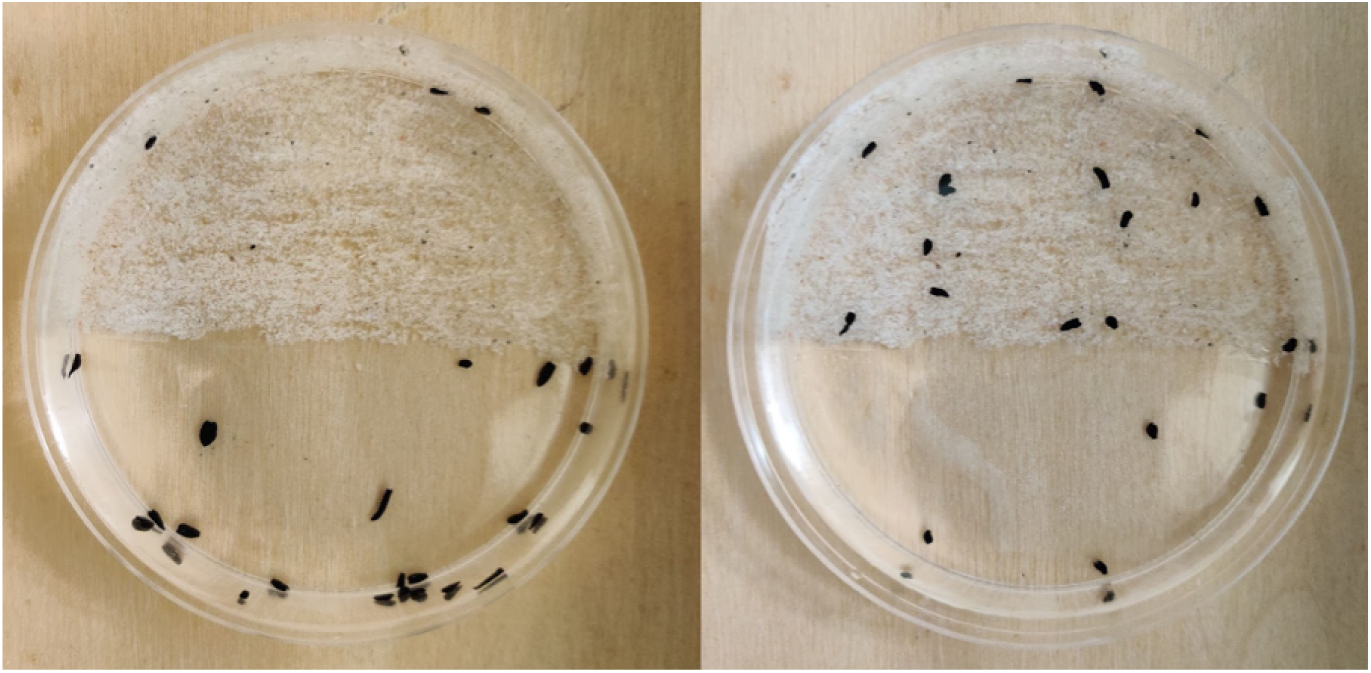
Left: Segments of the trained planaria on the half-rough dish after 30 minutes. Both the anterior and posterior segments show a strong preference for the smooth surface. Right: Segments of the untrained planaria on the half-rough dish after 30 minutes. The anterior as well as the posterior fragments prefer the rough surface.

**Figure 7:**
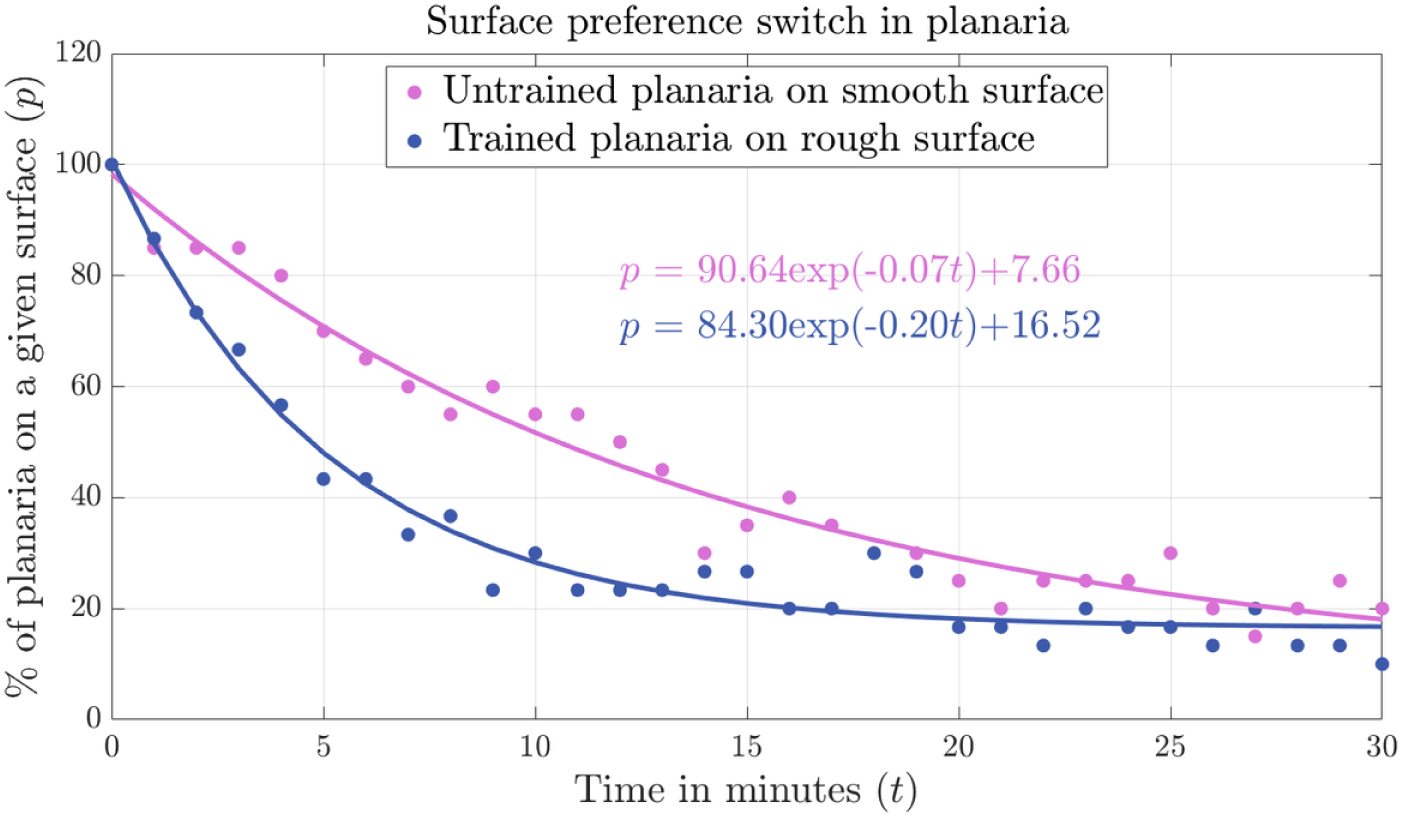
The switching of decapitated planaria between the rough and smooth surface, which is modeled by the exponential decay curves. The percentage of trained planaria segments present on the unpreferred rough surface is shown in blue and that of the untrained segments on the smooth (unpreferred) surface in purple. Both the groups tend to migrate toward the preferred surface gradually over 30 minutes. The trained planaria that were addicted to sugar exhibited a higher rate of transition toward the smooth surface, which might be associated with the influence of the addictive substance.

### 3.5 Dopamine Antagonist Treatment

The training phase consisted of alternating the planaria between the following surfaces and solutions for 12 days. The rough surface was paired with water (R+W) and the smooth surface was paired with a 10% sugar solution along with 1μM of dopamine antagonist (S+S+D). After the training phase, the surface preference of planaria was determined by observing the movement of each planaria on the half-rough dish for 30 minutes.

Fourteen out of 15 planaria did not show a preference shift as can be noticed from Fig. 8, meaning that they continued to prefer the rough surface while one switched the preference to the smooth surface.

**Figure 8:**
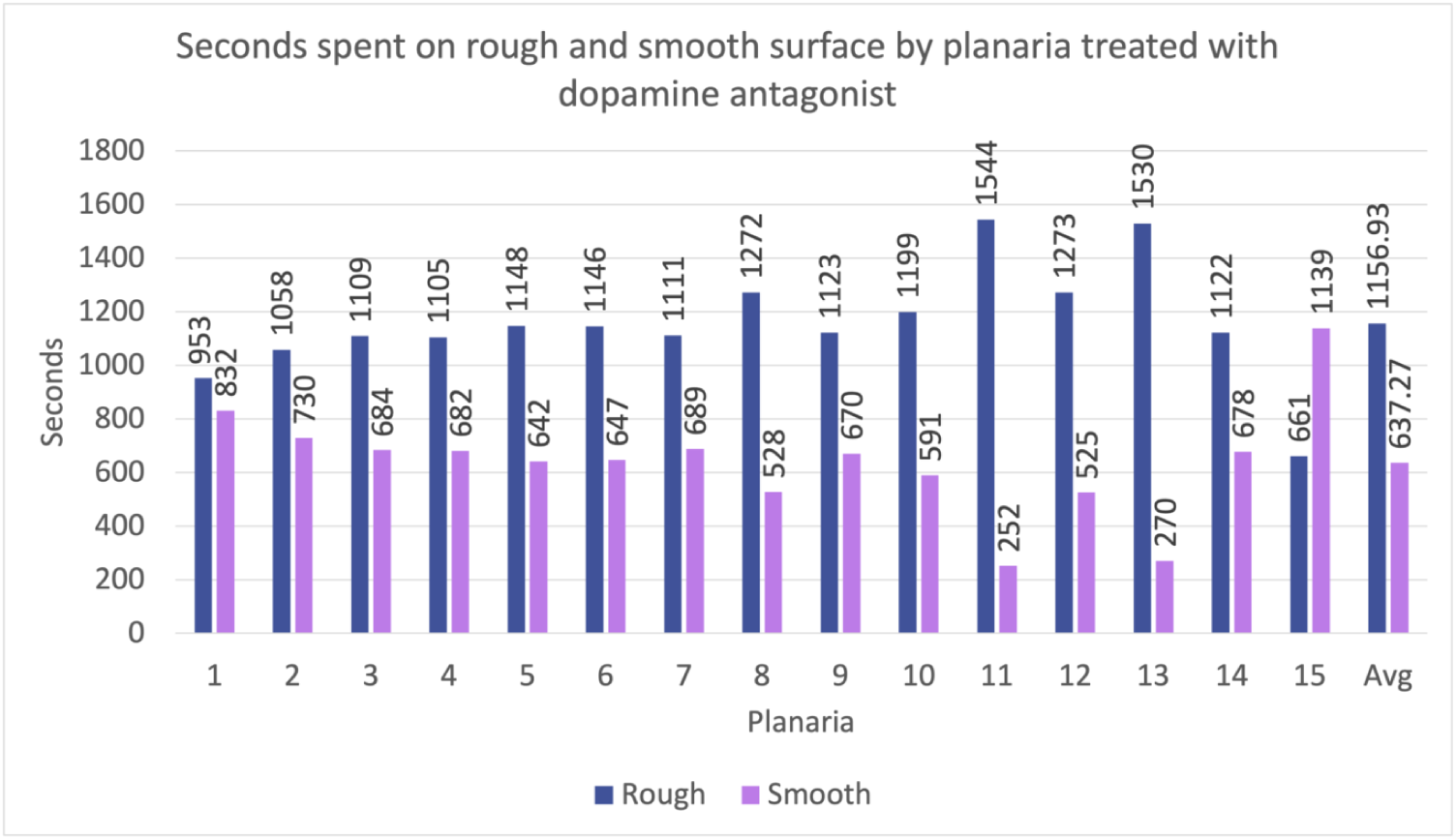
Surface preference of planaria trained with a dopamine antagonist. While 14 planaria did not shift their preference for the rough surface (blue), only one switched its preference to the smooth surface (purple).

### 3.6 Statistical Analysis

We performed the paired sample t-test to verify the statistical significance of our findings in the following scenarios: (i) the preference of untrained planaria for the rough surface (Section 3.2); (ii) the altered preference of trained planaria for the smooth surface (Section 3.3); and (iii) the preservation of surface preference in planaria due to the dopamine antagonist treatment (Section 3.5). In all the three scenarios, the inferences were found to be statistically significant at the 5% significance level, meaning that (i) the time spent by the untrained planaria on the rough surface is significantly longer than that on the smooth surface; (ii) the time spent by the trained planaria on the smooth surface is significantly longer than that on the rough surface; and (iii) the time spent by the dopamine-treated planaria on the rough surface is significantly longer than that on the smooth surface, thus preserving its inherent surface preference without being modified by the addictive substance. For the aforementioned cases, the *p-*values are reported in Table 1 and the box-and-whisker plots in Fig. 9 show the median, upper and lower quartile values, and the outliers.

**Table 1.** The *p*-value and statistical significance († ⇒ *p* < 0.05, †† ⇒ *p* < 0.01, † † † ⇒ *p* < 0.001, † † †† ⇒ *p* < 0.0001) of the difference between the time spent on the rough and smooth surface by the untrained, trained, and dopamine-treated planaria.

| Planaria Category | $p$-Value | Statistical Significance |
| --- | --- | --- |
| Untrained | 6.26e-06 | $\dagger\dagger\dagger\dagger$ |
| Trained | 1.88e-04 | $\dagger\dagger\dagger$ |
| Dopamine-Treated | 2.95e-04 | $\dagger\dagger\dagger$ |

**Figure 9:**
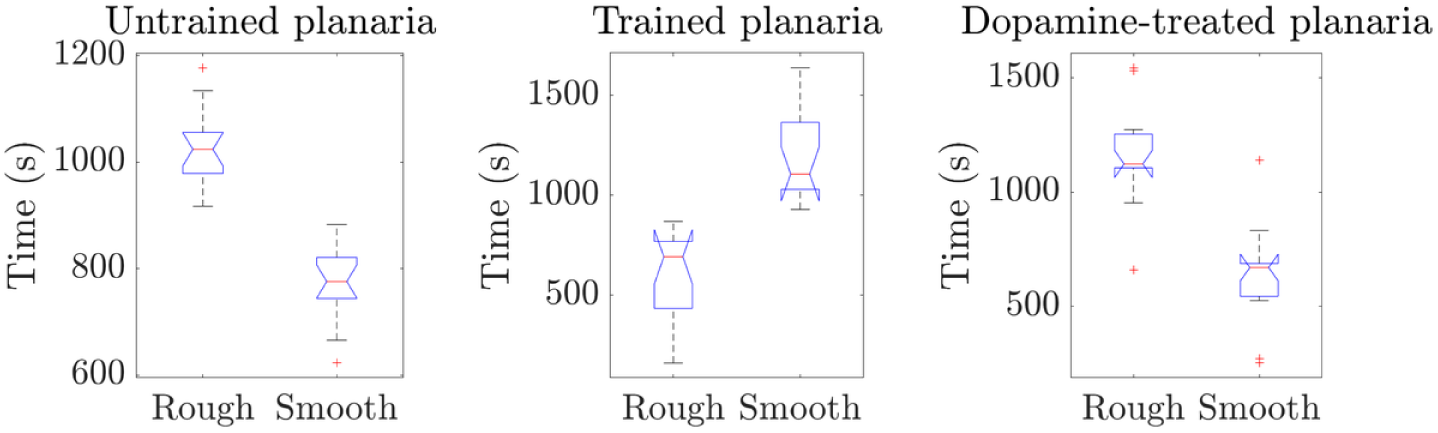
Box-and-whisker plots showing the median, upper and lower quartiles, and outliers of the time (in seconds) spent on the rough and smooth surface by the following categories of planaria: (left) untrained, (middle) trained, and (right) dopamine-treated. Notice that the interquartile ranges do not overlap in the boxplots, meaning that the difference between the time spent on the two surfaces is statistically significant in each scenario with the p-value reported in Table 1.

## 4 Discussions

### 4.1 Solution-Dependent Motility

The motility of planaria in the 10% sugar solution is noticeably higher than when they were placed in water. That is, the sucrose-induced motility measured by the number of gridline crossings is 208, whereas in water this measure is only 123. Presumably, the planaria would have attained an “excited state” due to the release of the neurotransmitter—dopamine—from the brain. This finding is in agreement with the outcome of the experiments recorded in (Pagán et al., 2009), where the planaria exposed to caffeine had increased motility compared to those exposed to dimethyl sulfoxide. It is due to the fact that caffeine is responsible for the release of extracellular glutamate and dopamine (Solinas et al., 2002). Our results, however, did not corroborate with the findings reported in (Jawad et al., 2017), where the planaria showed higher levels of activity in terms of the distance covered when placed in water than in the sucrose solution.

### 4.2 Establishment of CPP

The induction of a CPP was established in all the 15 planaria following the training for 12 days. While the untrained planaria showed a preference for the rough surface, their surface preference shifted to the smooth surface through Pavlovian conditioning (by pairing the rough surface with water and smooth surface with sugar). This shift in preference was consistent with the results obtained in (Jawad et al., 2017). However, unlike their approach, where they tracked the distance (in cm) covered by the planaria on either the rough or the smooth surface, we calculated the amount of time spent by them on each surface. A sucrose-induced CPP of planaria was realized in (Zhang et al., 2013); but instead of pairing different surfaces with solutions, they paired the sugar solution with light (unpreferred environment) and water with darkness (preferred environment).

All the untrained planaria showed a strong preference for the rough surface, as evidenced by the average amount of time spent on the rough surface (1025.53 seconds) as opposed to (774.47 seconds) on the smooth surface. It suggests that the unconditioned organisms have an innate preference for the rough surface. After training, a major duration of the 30-minute interval was spent on the smooth surface of the half-rough dish (an average of 1193 seconds on the smooth surface in contrast to just 607 seconds on the rough surface). Hence, all the trained planaria preferred the smooth surface as they associated it with the sucrose solution. This verifies that a CPP had been established, meaning that addiction to sucrose had been instilled.

### 4.3 Effect of Sucrose Concentration & Exposure Duration on CPP

The rationale for fixing the concentration of sugar solution at 10% for all the experiments is stated below. Sucrose concentrations between 0.1% and 10% have been shown to have the greatest effect on conditioning (Zhang et al., 2013). Even though the concentrations less than 0.1% and more than 10% happen to produce apreference shift in planaria, the effect was shown to be statistically insignificant (Zhang et al., 2013). The first phase of training was conducted by alternating between the R+W (rough+water) and S+S (smooth+sugar solution) every 24 hours. This did not produce any significant result, which might be due to overexposure and withdrawal effects as remarked in (Sacavage et al., 2008). The second training phase comprised shorter exposure durations (30 minutes per day) since longer exposure periods such as a day would cause withdrawal effects once the planaria were out of the addictive solution (Sacavage et al., 2008). Similar withdrawal effects were noticed in planaria exposed to a sugar solution for an hour every day in the investigation carried out in (Zhang et al., 2013).

### 4.4 Addiction-Related Memory Retention

Planaria are known to learn and retain memory. Long-term memory retention has been reported in previous studies (Shomrat and Levin, 2013), where a planaria’ s memory of the environment seemed to persist for more than 14 days. In the case of a CPP, it has been shown in (Jawad et al., 2017) that planaria could retain addictive memory for four days after the training phase until they reached the extinction phase, during when the unpreferred surface is no longer associated with the sugar environment. Nevertheless, in order to proceed with caution in our experiment, the planaria were trained for an additional duration of four days after the post-training phase (when the addictive behavior of the planaria was observed in the half-rough dish) before they were decapitated and tested for memory retention.

### 4.5 Addictive Memory Transfer and Possible Implications

It takes about two weeks for an amputated planaria to undergo complete regeneration and form into two separate organisms (Hammoudi et al., 2018). In a previous study performed on memory retention in planaria, it has been demonstrated that memory of the familiarized environment can be retrieved by decapitated planaria once their heads regenerated (Shomrat and Levin, 2013). Their experimental findings suggest that the trained planaria could remember the feeding procedure and have a shorter feeding latency than the untrained ones (Shomrat and Levin, 2013).

Understanding the mechanisms that govern the memory transfer of sugar addiction in planaria regenerated from amputated posterior segments can provide valuable pointers to prevent addictions of humans to other substances as well. Sucrose is shown to have the same effects as some addictive substances and evoke the same reactions in the brain (Jawad et al., 2017). Drug abuse has been shown to increase the amount of extracellular dopamine, which mimics the physiological dopamine produced (Volkow et al., 2009). Interestingly, sugar has been proven to increase the amount of dopamine and opioid released and engages the same pathways that are activated by addictive drugs (Avena et al., 2008).

### 4.6 Prospects of Dopamine Antagonist to Treat Addictions

The role of the antagonist lies with its fixation to the receptors, thus preventing the dopamine from fixing to the dopamine receptors. While adding 1μM of D1 dopamine antagonist to the sugar solution, 14 out of 15 planaria failed to exhibit any CPP and preferred the rough surface instead (an average of 1156.93 seconds on the rough surface and 637.27 seconds on the smooth surface). This is consistent with the results reported in (Jawad et al., 2017) and (Zhang et al., 2013). A lack of CPP formation implies that the addiction of the planaria gets disrupted. Since the dopamine antagonist plays a preventive role in the development of sucrose addiction, this knowledge can be translated into devising mechanisms to disrupt other addictions to substance abuse.

### 4.7 Limitations and Future Perspectives

Admittedly, there are some inherent limitations in the experimental procedure. The sample size of 15 chosen for every experiment may not be sufficient enough to arrive at concrete conclusions. Therefore, future research should include a greater number of specimens in order to verify the hypotheses with statistical tests. After experimenting with both the black planaria (*Dugesia dorotocephala*) and the brown ones (*Dugesia tigrina*), we realized that the black planaria were seemingly more prone to acquire the sugar-related addiction and hence would be suitable for the experiments. Computerized movement detection of planaria would be more accurate to account for the time spent on different surfaces and would evade possible human errors. The secondary hypothesis may further be validated by treating the planaria with D2, D3, D4, and D5 dopamine antagonists. Further research can be carried out to investigate whether memory transfer in planaria occurs at the genetic level through a molecular approach.

